# Reducing structural non-identifiabilities in upstream bioprocess models using profile-likelihood

**DOI:** 10.1101/2022.02.17.480405

**Authors:** Heiko Babel, Ola Omar, Albert Paul, Joachim Bär

## Abstract

Process models are increasingly used to support upstream process development in the biopharmaceutical industry for process optimization, scale-up and to reduce experimental effort. Parametric unstructured models based biological mechanisms are highly promising, since they do not require large amounts of data. The critical part in the application is the certainty of the parameter estimates, since uncertainty of the parameter estimates propagates to model predictions and can increase the risk associated with those predictions. Currently Fisher-Information-Matrix based approximations or Monte-Carlo approaches are used to estimate parameter confidence intervals and regularization approaches to decrease parameter uncertainty.

Here we apply profile likelihood to determine parameter identifiability of a recent upstream process model. We have investigated the effect of data amount on identifiability and found out that addition of data reduces non-identifiability. The likelihood profiles of non-identifiable parameters were then used to uncover structural model changes. These changes effectively alleviate the remaining non-identifiabilities except for a single parameter out of 21 total parameters.

We present the first application of profile likelihood to a complete upstream process model. Profile likelihood is a highly suitable method to determine parameter confidence intervals in upstream process models and provides reliable estimates even with non-linear models and limited data.

## Introduction

Mathematical models find increasingly application in biopharmaceutical process development, which is encouraged by regulatory agencies through the PAT initiative of the FDA (FDA, 2004) and recommended for the use of process validation (FDA, 2011). In upstream processing mathematical models should increase the process understanding, reduce the experimental effort, increase the productivity and facilitate a Quality-by-Design approach (Narayanan et al., 2019; Varsakelis et al., 2020; Zobel-Roos et al., 2019). Currently there are different approaches used to model upstream bioprocesses. So-called hybrid models combining machine learning and mechanistic parts show promising results (Narayanan et al., 2019; Popp et al., 2016) for process optimization and process understanding, however parametrization or training of such hybrid models requires either larger amounts of data or expensive offline analytics of metabolites with a high time resolution.

In contrast macroscopic unstructured models use mechanistic insight and can be parametrized with single fermentation experiments. Unstructured models have been used for different approaches. Promising applications are process optimization (Möller et al., 2019a) and optimization of pH shifts (Paul et al., 2019). They have been used to uncover biological mechanisms that affect scale-up (Möller et al., 2019b), but also other upstream processes such as the seed-train have successfully been optimized with unstructured models (Hernández Rodríguez et al., 2019; Kern et al., 2016).

One major challenge in the use of macroscopic models is the determination of the model parameters. With non-linear models and limited data availability model parameters can be difficult to determine (Raue et al., 2009; Ulonska et al., 2018). Consequently the uncertainty in the parameter estimates propagates into uncertainty of the simulation (Anane et al., 2019; Kreutz et al., 2012; Liu and Gunawan, 2017b; Maiwald et al., 2016). It is therefore of high importance to determine not only the parameter values but also the confidence intervals of the parameters in a reliable manner, since this can critically affect the risk associated with the model predictions.

To this end several methods have been used (Rajamanickam et al., 2021). The classical approach uses numerical estimates of the Jacobi Matrix and approximates the confidence interval with a parabola (Meeker and Escobar, 1995), also known as Covariance approach or Fisher-Information Matrix (FIM) (Rischawy et al., 2019; Ulonska et al., 2018; Wieland et al., 2021). This estimate is true under the assumption of normal distributed errors and large number of data (Neale and Miller, 1997; Seber and Wild, 2003). This method is known to over or underestimate true confidence intervals non-linear models with limited data (Moerbeek et al., 2004; Wieland et al., 2021). The method also relies on the numerical estimation of sensitivities or the Jacobi matrix which can be error-prone (Wieland et al., 2021). Another approach are Monte-Carlo approaches (Liu and Gunawan, 2017b; Möller et al., 2019b) which have the problem of handling unknown covariances and fine tuning of the sampling algorithm without autocorrelation (Hug et al., 2013).

In the field of systems biology an alternative approach to parameter confidence intervals has been established, the so-called profile-likelihood (Raue et al., 2009). In contrast to the FIM approach the profile-likelihood is an exact method and results in non-symmetric confidence intervals. Profile-likelihood is suited for non-linear models that are parametrized with a limited number of data-points (Moerbeek et al., 2004; Neale and Miller, 1997). In bioprocess modeling this approach has only been applied once by Kroll et. al. (Kroll et al., 2017). Here a model only for the viable cell density was parametrized with a single run with high quality data and high time resolution. It is currently unknown if the profile-likelihood method can be applied to complete upstream process models comprising all relevant state-variables, i.e. not only viable cell concentration but also dead cell, product and metabolite concentrations. To consider all available concentrations is of importance since metabolite concentrations and viability also influence product quality (Paul et al., 2018; Xu et al., 2016). Compared to Kroll et. al. (Kroll et al., 2017) process data is usually sampled daily with less precision. So far parameters of complete upstream processes have only been characterized by approximate confidence intervals but not exact confidence intervals (Ulonska et al., 2018) and exact confidence intervals for such process models with typical process data have not been determined.

To determine exact confidence intervals of complete process models we use the process model of Ulonska et. al. (Ulonska et al., 2018) comprising viable cell, dead cell, glucose, glutamine, lactate, ammonia and mAb product concentration and thereby modeling relevant process parameters. We then studied the effect of data availability by comparing profile likelihoods obtained from a single run and a data set with differential feed rate and seeding cell density. Next, we employed a profile-likelihood guided analysis to change the model structure and alleviated remaining non-identifiabilities. Finally, we compared the different model structures with increased parameter identifiability with respect to their fit to the experimental data.

## Material and Methods

### Experimental Data

We used the experimental data from the fed-batch cultivation of a CHO-DG44 cell line cultivated in a chemically defined medium. The five fermentation experiments to produce a monoclonal antibody were all conducted in controlled automated miniaturized bioreactor (ambr250) systems (Sartorius AG, Germany) for 14 days. Beside the center point, the data of additional four runs varying in feed rate and seeding cell density were used. Compared to the ‘center point’ the seeding cell density was increased or decreased by 30 %. The feed rate in the were decreased by 25 % and increased by 12 % respectively compared to the ‘center point’. The feed was started after 24h of the cultivation start time. Apart from these variations, the cultivation took place under the same conditions. The temperature and dissolved oxygen was held constant. Starting from day 2, glucose boluses were applied and until day 3 glutamine boluses were added.

The glucose, glutamine, lactate and ammonia concentrations were measured daily using the Prime60 system (Software: Konelab Prime 7.2.1, Co. Thermo Fisher Scientific, USA). The titer concentration was measured starting from day 8 via IGG method using the Prime60 system. The viable cell density and the total cell density were measured using the Cedex HiRes (Roche, Switzerland). Dead cell density was calculated from the difference between viable and total cell density.

### Mathematical Model

The process model was adapted from Ulonska et. al. (Ulonska et al., 2018) and is described in detail there. The differential equations and kinetics are shown in the supplementary text S1. In brief the model contains viable cell density, dead cells (or total cell density) and cell lysis. Product formation is proportional to viable cell concentration. Cell growth depends on glucose and glutamine and a glutamine containing supplement. In addition, the metabolites lactate and ammonia are considered. The kinetics were modeled using zero and first-order rates, Monod-terms and rational inhibition terms.

### Model Parametrization and Profile Likelihood Analysis

We use the weighted sum of squared residuals as objective function for the parametrization of the models:

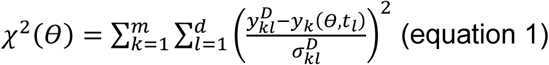

here y_kl_^D^ is the data-point of the k-th observable at the l-th measurement timepoint, y(θ,t_l_) is the model prediction for the corresponding datapoint with the parameter set θ. σ_kl_^D^ is the standard deviation of the measurement. Under the assumption of normal distributed observational noise the weighted sum of squared residuals corresponds to the maximum likelihood estimate (MLE) (Raue et al., 2009), since then *χ*^2^(*θ*) = *const* − 2*Log*(*L*(*θ*)).

Instead of ‘normal theory’ based confidence regions derived from the asymptotic distribution of the ‘Wald statistic’ we use parameter confidence intervals derived from likelihood ratio tests (Meeker and Escobar, 1995):

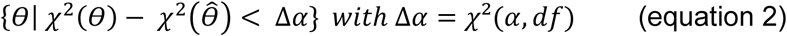

Wherein the borders represent the confidence interval with confidence level α. The threshold Δ_α_ is the α quantile of the Χ^2^-distribution. When df=1 the pointwise confidence interval are obtained, and df = #θ give simultaneous confidence levels (Raue et al., 2009). For finite samples likelihood based confidence intervals are considered superior compared to the asymptotic distribution of the ‘Wald statistic’ (Moerbeek et al., 2004; Neale and Miller, 1997).

Parameter identifiability is determined by the profile likelihood given by:

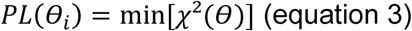

A parameter is classified as ‘identifiable’ if the profile likelihood reaches the confidence threshold for lower and higher parameter values then the optimal parameter value.

The normalized root-mean squared error was calculated for each observable by:

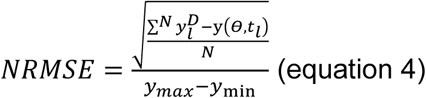

All calculations were performed in Matlab R2019a and profile-likelihoods were calculated using the data2dynamics toolbox (Raue et al., 2015) with the lsqnonlin optimizer of Matlab. Initial model parameters were estimated using a global optimization approach with latin-hypercube sampling within specified parameter bounds. Convergence was checked using the arLocalLHS function of data2dynamics.

## Results

### Parameter identifiability depends on the data

The minimal dataset for an upstream process models consists of a single fermentation. We fitted the state-of-the-art process model described by Ulonska et. al. (Ulonska et al., 2018) to a single center point run of a process development study (black curve in Figure 1). The model captures the dynamics of cell-growth and death (VCC and DCC) well, going from growth to death phase. The model fit agrees with the experimental data of glucose, lactate and glutamine and captures the product formation. It performs poorly for the ammonia formation. The experimental data showed for ammonia an initial increase until day 3, a slight decrease and then further increase starting at day 7 until the end of the process. Here, the model only increased ammonia values until day 4 and then stays constant with a slight decline until the end of the fermentation.

**Figure 1:**
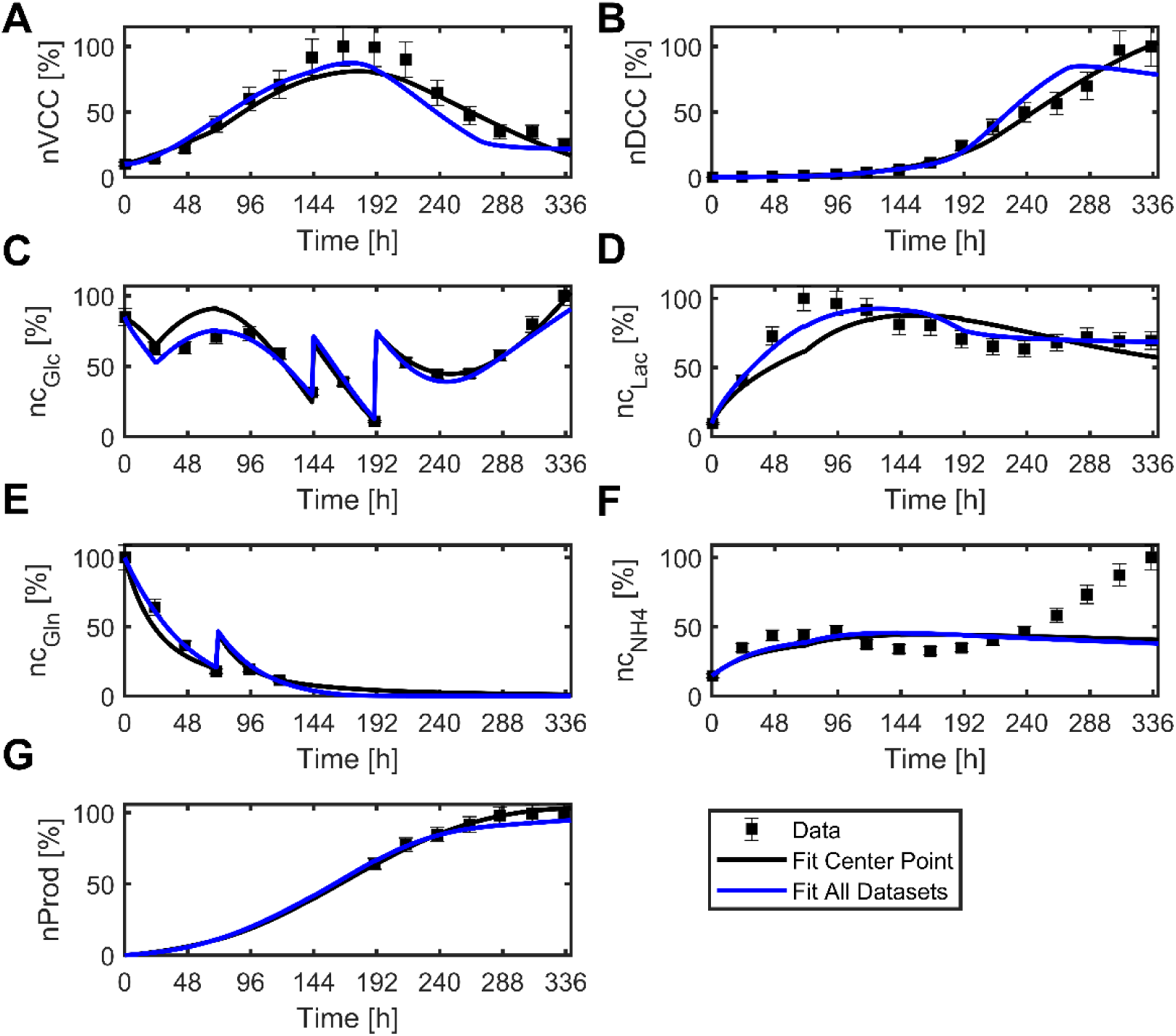
Original model fit to center point or full dataset. All curves are normalized to the maximal data-value. Experimental data is marked with square-symbols and the error-bar correspond to the standard deviation from the analytical measurement method. Lines denote the model simulation of the center-point run of a model parametrized only with the center-point (black line) or with the center point and 4 additional runs (blue line).

We next assessed if the model parameters that were determined from the single center point run are identifiable. The identifiability was determined by calculation of the profile-likelihood for each fit parameter. Of the 21 parameters 8 parameters could not be identified with the single run. Figure 2 shows that several profile likelihoods of the parametrization with a single run (black lines) do not hit the 95 % confidence threshold and do not significantly increase in the observed parameter range. The parameters biomass yields from glucose and glutamine, Y_BM,Glc_ and Y_BM,Gln_ respectively, have both a lower bound but no upper boundary. For the Monod-constant of lactate uptake k_Lac_, the maximal Lactate uptake rate q_Lac,m_ and the glucose maintenance m_Glc_ we do not observe any curvature in the profile likelihood, but only a flat profile. The inhibition constants ν_Lac,Gln_ and ν_Lac,Prod_ have no lower and upper boundary, respectively. The maximal glutamine uptake rate q_Gln,m_ appears to mirror the biomass/glutamine yield coefficient with a defined minimum but no lower boundary. In the cases of e.g. the biomass/glutamine yield Y_BM,Gln_ but also the ammonia formation rate r_NH4,Gln_ the approximation with a parabola significantly underestimates the true parameter confidence interval (supplementary table 1).

**Figure 2:**
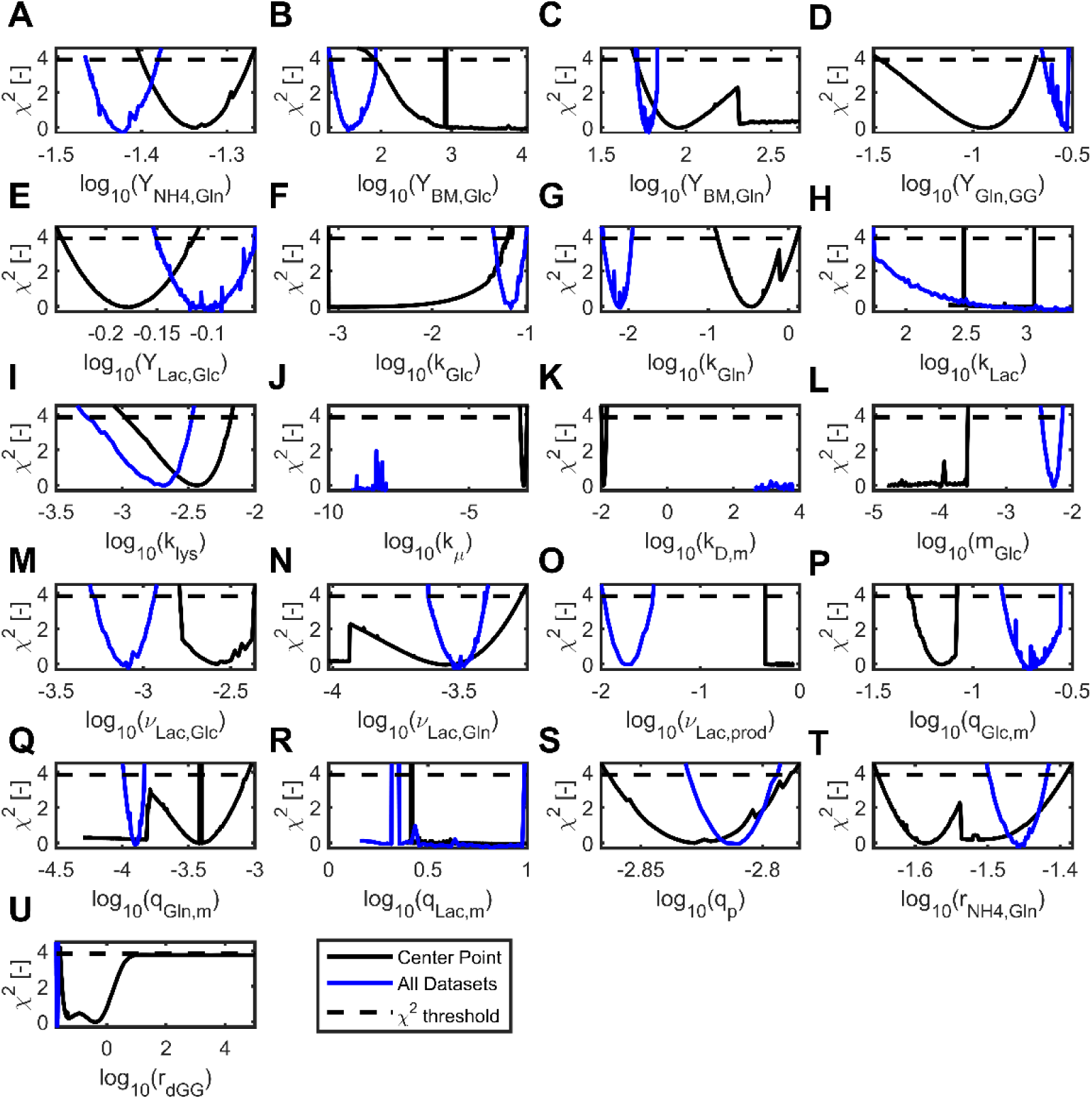
Profile-Likelihood of parameters from a model fit to center point and full dataset. Profile likelihood from the model parametrized with a single center point run are shown in black, the model parametrized with the full dataset is shown in blue. The dashed line shows the point-wise 95 % confidence level. The minimum value of the objective function Χ^2^ for each parametrization was subtracted from the profiles for comparability

Next, we wondered if these non-identifiabilities could be alleviated by parametrization of the model with additional datasets. To this end four additional runs with variations in seeding-cell density and feed rate were added and the model was reparametrized. The reparametrized model does not show differences in its simulation of the center point (blue line in Figure 1). We observe small deviations at day 8 compared to the previous parametrization in VCC and DCC. Also, the simulation for DCC deviates from the data at the final days of the process. The simulation of the lactate concentration is slightly shifted in the reparametrized model and the lactate peak in the data appears to correspond better with the simulation.

As shown in figure 2 most of the optimal parameter values and therefore also the profile likelihoods are shifted by the addition of 4 complementary runs. For the two biomass yield coefficients Y_BM,Glc_ and Y_BM,Gln_, the inhibition constants ν_Lac,Gln_ and ν_Lac,Prod_, the glucose maintenance m_Glc_ and maximal Glutamine uptake rate q_Gln,m_ the non-identifiabilities were resolved by the addition of new data, since the likelihood profiles now hit the limit for the 95 % confidence interval. The non-identifiability of the lactate uptake parameters k_Lac_ and q_Lac,m_ could not be resolved by the additional experiments. Unexpectedly the addition of new data introduced new non-identifiability in the cell death kinetic. Both parameters of the cell-death kinetic, the maximal cell death rate k_D,m_ and the inhibition constant by cell growth k_μ_ cannot be identified anymore. Data addition alleviated most non-identifiabilities, however some non-identifiabilities remained.

### Structural non-identifiability of Lactate uptake kinetic and cell death kinetic

We next examined the remaining non-identifiabilities in detail. To this end we plotted the observed parameter changes in the profile-likelihood calculation of k_Lac_ and q_Lac,m_ of the lactate kinetic and k_D,m_ and k_μ_ of the cell death kinetic (Figure 3). For all those profiles the X^2^ remained below 95 % confidence threshold, except for k_Lac_ where the profile reaches the 95 % confidence threshold for lower values then the optimal value.

**Figure 3:**
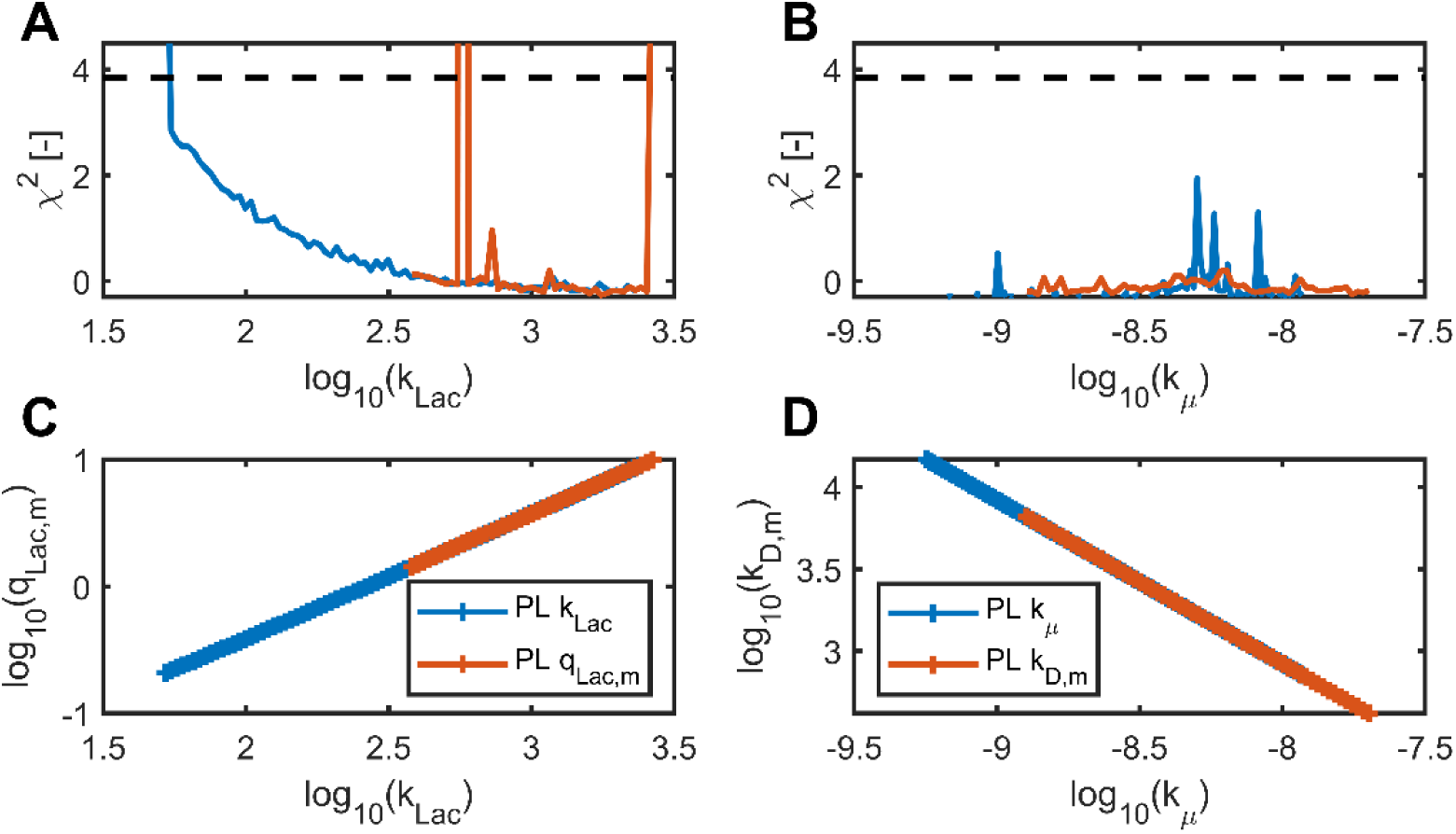
Non-Identifiabilities in model with full dataset. (A-B) Profile likelihood of lactate uptake kinetic parameters and cell death kinetic parameters as a function of k_Lac_ and k_μ_ respectively. Dashed line shows 95 % confidence threshold. (C) Optimized parameter values of q_Lac,m_ and k_Lac_ of the profile-likelihoods for q_Lac,m_ and k_Lac_. (D) Optimized parameter values of k_D,m_ and k_μ_ of the profile-likelihoods for k_D,m_ and k_μ_.

Interestingly, for all profile-likelihood calculations we observe a linear correlation of the two parameter combinations. In the case of the lactate kinetic, k_Lac_ and q_Lac,m_ exhibit a positive linear correlation with a slope of 0.99. The relation we observe is approximately log_10_ *q*_*Lac,m*_ ≈ log_10_ *k*_*Lac*_ + *C* which means that:

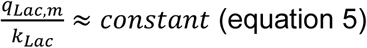

The range of optimal parameter values for k_Lac_ in the parametrization extends from 10^2^ to 10^3.5^ g/L, this is highly above the observed lactate concentrations in CHO based bioprocesses. In this case the Monod-term in the lactate uptake kinetic can be approximated with the limit of c_Lac_<<k_Lac_ and becomes:

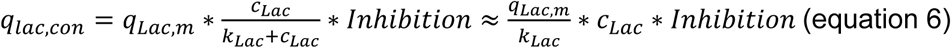

To alleviate this non-identifiability the lactate uptake kinetic could be replaced by this approximation. This yields a new effective parameter 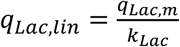 that replaces q_Lac,m_ and k_Lac_ in the model.

For the cell death kinetic the correlation between k_D,m_ and k_μ_ is negative with a slope of -0.99. In contrast to the lactate uptake parameters here the product of k_D,m_ and k_μ_ is constant:

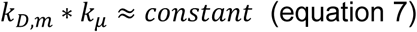

In this parametrization the values of k_μ_ are very small and the cell death kinetic can be approximated for the limit that k_μ_ << μ to:

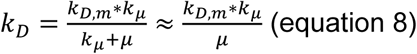

Here it is not advisable to replace the current term with the approximated version since this could results in divisions by zero, however to alleviate the non-identifiability of k_D,m_ we suggest to remove k_μ_ from the numerator.

### Model structure adaptation alleviate non-identifiability

Based on the profile-likelihood calculations we could identify structural changes in the model that might alleviate the observed parameter non-identifiabilities. In a next step we implemented those structural model changes to see if they would indeed result in identifiable parameter values. We implemented single structural changes first and then combined them together in a fully adapted model.

As shown in figure 4 the remaining lactate-kinetic parameter q_Lac,lin_ is identifiable in the model where only the lactate uptake kinetic is changed and the fully adapted model. Similarly we observe that the death rate parameter k_D,m_ can be identified in the model where only the cell death rate is adapted and the fully adapted model. As expected, the cell death parameter k_μ_ can still not be identified as it tends to 0. We observe that the optimal parameter values are shifted slightly to higher values in the fully adapted model compared to the models with single model term adaptations.

**Figure 4:**
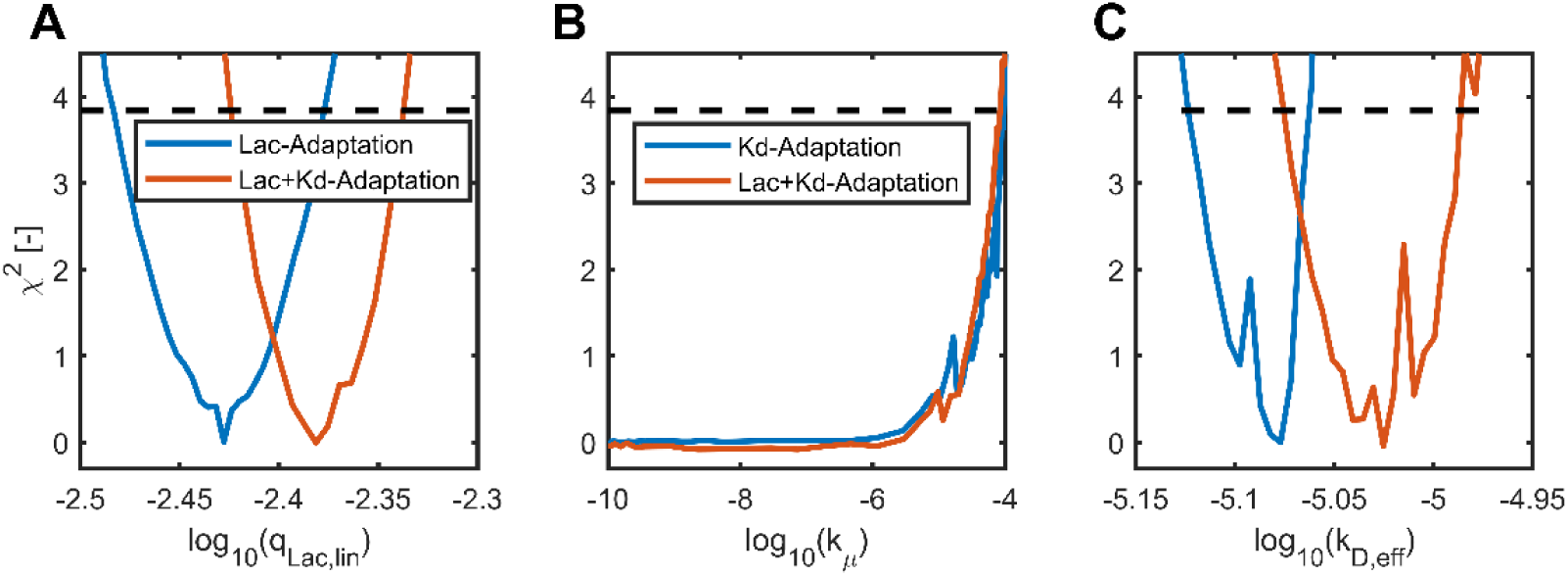
Profile Likelihood of Parameters of Adapted Models (A) Profile likelihood of q_Lac,lin_ for a model where only the lactate kinetic is adapted and both lactate and cell death kinetic are adapted. (B) Profile likelihood of k_μ_ for a model where only the cell death kinetic is adapted and both lactate and cell death kinetic are adapted. (C) Profile likelihood of k_D,m_ for a model where only the cell death kinetic is adapted and both lactate and cell death kinetic are adapted.

Finally, we wanted to know how those adaptations in the model structure affect the overall fit of the model. To this end we calculated the normalized root mean square error (NRMSE) for all observables and the different models. We observe only slight differences in the model performance for all single model adaptations compared to the original model (figure 5). The strongest deviations can be found for the fully adapted model. Here the NRMSE is increased by a value of 0.01 for both the dead cell concentration DCC and product concentration and increased by a value of 0.003 for VCC. However, the error is decreased by a value of 0.01 for both glucose and lactate concentration and decreased by values of 0.004 and 0.001 for glutamine and ammonia concentration respectively.

**Figure 5:**
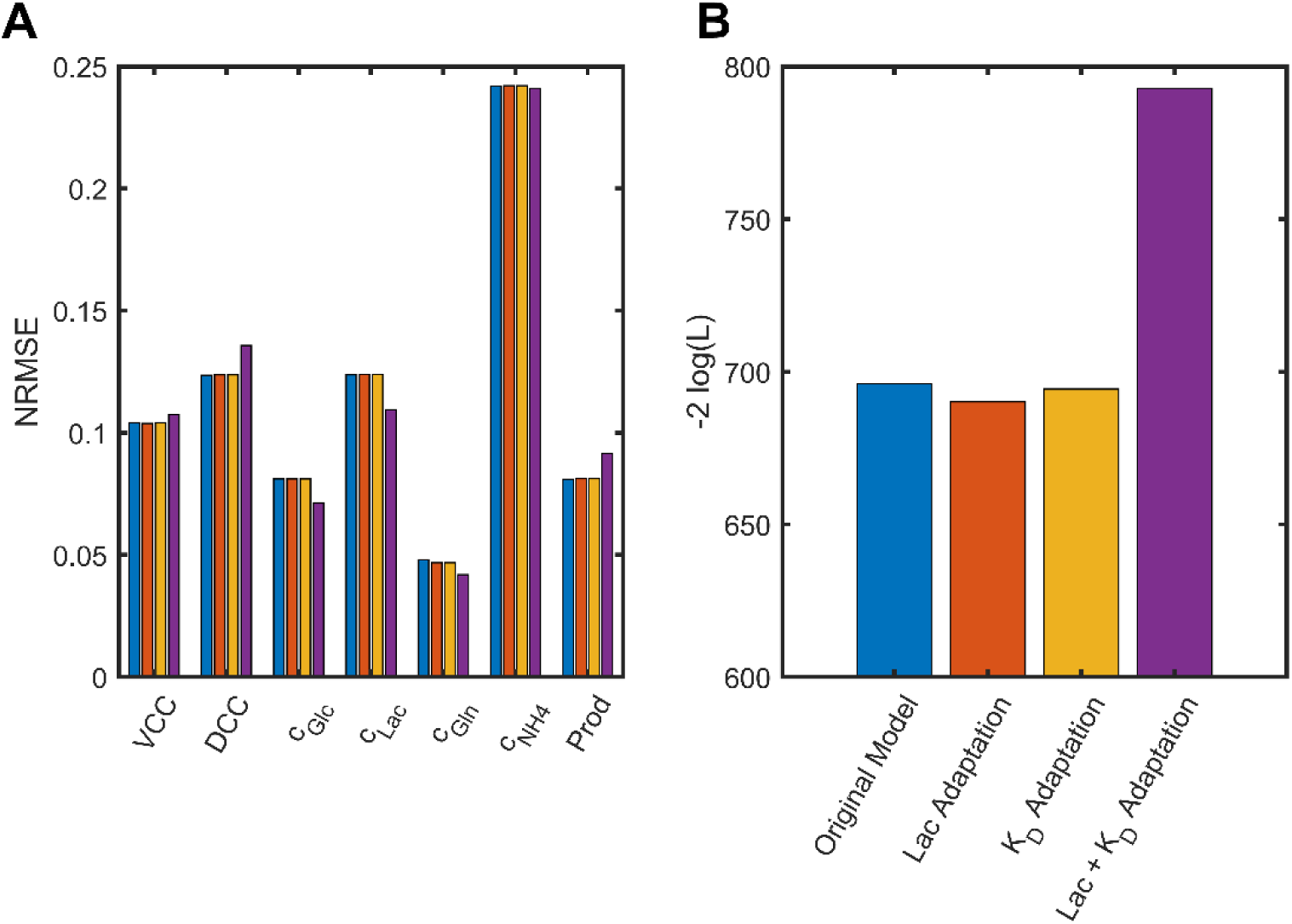
Model fit of different model adaptations (A) Normalized root mean squared error (NRMSE) for the observables with the original model, the lactate kinetic adapted model, the cell death kinetic adapted model and the fully adapted model. (B) Negative Log-Likelihood of the different models.

A different picture emerges if we look at the negative Log-Likelihood value of each model, which can be derived from the objective function (equation 1). The negative log-likelihood decreases when the lactate kinetic or the cell death kinetic are adapted by values of -2 log(L) of 5.9 and 1.7, respectively. However, when both adaptations are implemented in the model, the negative log-likelihood strongly increases by a -2 log(L) value of 97.

## Discussion

The process model used in this work and previously described by Ulonska et. al. (Ulonska et al., 2018) describes fed-batch fermentation processes accurately, which we could confirm in this study with new data. Ulonska et. al. also described only a reasonable model fit to the ammonia data. The simulations showed an increase in ammonia concentration and then a plateau, however in the data clearly two periods of concentration increases can be observed. In this macroscopic model glutamine and a glutamine containing supplement are considered as sources for ammonia. Defined media used in current fed-batch processes also contain amino acids in the feed medium (Xing et al., 2011). Amino acids can also be a source for ammonia but are not described in this model. Addition of these additional ammonia sources might increase the descriptive capability of this model.

We could show that 6 parameters could be identified after the addition of new runs. The additional runs had variations in seeding cell density and in feed rate. Both experiments effectively change the nutrient availability. Variations in feed-rate change the available glucose that is provided in the feed, which can lead to the identification of parameters associated with glucose consumption, i.e. the yield and the maintenance metabolism. When the seeding cell density is shifted, this changes both the glutamine and glucose nutrient availability. Consequently, the yield coefficient for biomass from glutamine and the maximal glutamine uptake rate can be identified. The changes in nutrient availability also triggers changes in the metabolism, and therefore additional variations in lactate concentrations are introduced with the new dataset which enable the identification of the inhibition constants of the lactate production and consumption.

Since we used the same model as Ulonska et. al. (Ulonska et al., 2018) we can directly compare the estimated parameter values. 9 of the 19 parameter values published there are in the same range. There are three major differences. Due to the non-identifiabilities determined in our study we have deviating parameter values for the lactate uptake parameters and the cell-death kinetic. This is to be expected since non-identifiabilities result in large confidence intervals for the parameter values. The biomass yield for glutamine and glucose have higher parameter values in our model, this is however compensated by lower uptake rates. Finally, our data suggests a higher production rate. The deviating parameter values are highly dependent on the clone and cell line (Ulonska et al., 2018) but also the process scale (Möller et al., 2019b) and media composition (Robitaille et al., 2015) can affect those parameters.

Although data addition alleviated parameter identifiabilities, one non-identifiability persisted, and an additional non-identifiability occurred. The lactate-uptake kinetic parameters could not be identified with the available dataset. The likelihood-profile showed a distinct lower limit of k_Lac_ making it a practical non-identifiability. In the used dataset no lactate concentrations are in the concentration range of k_Lac_ where the optimal parameter value is 10^3^ g/L. In principle this non-identifiability could be removed by adding data where cells are treated with lactate concentrations >10^3^ g/L. However, this is not feasible due to practical reasons such as the solubility of lactate and that additional effects might be observed by high lactate concentrations like growth inhibition (Lao and Toth, 1997). Therefore, the model structure needs to be adapted. Here we replaced the Michaelis-Menten uptake kinetic with its approximation for small substrate concentrations, i.e. a linear lactate uptake rate. We could reduce the number of model parameters from 21 to 20 and the new effective parameter q_Lac,lin_ could successfully be identified from the available dataset. Moreover, introduction of this model structure change increased the fit significantly (ΔΧ^2^=5.98 p=0.0145 with df=1). These types of non-identifiabilities are a common problem in biological models and readily resolved after identification with the profile likelihood (Maiwald et al., 2016).

A more complex situation arose for the cell death kinetic. Here, we observed identifiability of the cell death parameters for the single center point run, but the parameters became non-identifiable after additional runs were added. We also observed that the model fit to the center point becomes worse if more data is added at the end of the process for the dead cell concentration (Figure 1 B). The profile-likelihood analysis revealed that the product of k_D,m_ and k_μ_ is in fact constant and replacing this product in the cell-death kinetic with a new parameter rendered the new parameter identifiable. But the non-identifiability of k_μ_ could not be resolved. We suggest that the non-identifiability arose due to a biological phenomenon that is observed in the data but not covered by the model structure. The experimental data in figure 1 showed that the viable cell concentration is constant after 288 hours, at the same time the dead cell concentration increases. Ignoring dilution effects, the viable cell concentration is described by:

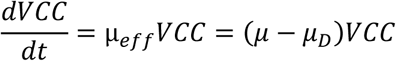

since we observe a positive death rate (increasing DCC) μ_D_>0 and stationary viable cell concentration μ_eff_=0 it follows that cells still replicate although they are at the same time dying and that μ>0. The cell death kinetic is formulated such that cell growth μ>0 and cell death μ_D_>0 are mutually exclusive. Here we suggest that for our dataset a different mechanism plays a role that allows for this type of stationary viable cell concentration. Cell death might depend on limiting substrate concentrations or accumulation of toxic compounds as e.g. in proposed in the model of Pörtner et. al. (Möller et al., 2019b).

The likelihood profiles obtained in this study are often skewed and show the non-linearity of the parameter estimation problem for upstream process models. Especially for model parametrization with only one control run the profiles initially resemble parabolas but then have abrupt jumps. Jumps in profile-likelihood have been observed previously (Raue et al., 2009) and originate from local likelihood minima that are in close proximity to the global optimum. Here, approaches based on the Fisher-Information Matrix (FIM) or covariance (Ulonska et al., 2018), i.e. based on the Jacobi matrix at the optimal parameter point strongly over or underestimate the true parameter confidence intervals (see supplementary table 1). Figure 2 shows some symmetric confidence intervals; however, the parameter values are plotted on a log scale. Therefore, an exponential transformation would reveal that most parameters confidence intervals indeed do have asymmetric confidence interval on a linear scale. Methods which assume symmetric confidence intervals such as the FIM will in this case still over or underestimate the true confidence intervals. The likelihood profiles calculated from the full dataset are rather noisy. This can be due to the roughness of the optimization problem at hand, which might be introduced by non-steady events such as feed-starts and bolus addition in the simulation. Also, the profile-likelihood depends critically on the optimizer used. Only if the optimizer reaches an optimum a valid profile-likelihood can be obtained. Here adjustment of the optimizer parameters or different optimizers might reduce the noise currently observed in the likelihood profiles.

The uncertainty in the data of upstream bioprocesses and in the predictions of models have been identified as a critical issue in seed train predictions (Hernández Rodríguez et al., 2019), scale-up prediction (Möller et al., 2019b) and process optimization (Liu and Gunawan, 2017a). It has also been shown previously that uncertainty in the model parameters propagates to uncertainty of model predictions (Anane et al., 2019). Since uncertain predictions of process performance are from little value, only models with small uncertainties are of value for process development and commercial model use. It is therefore of high interest to correctly quantify parameter uncertainty and potentially decrease uncertainty of parameters. The profile-likelihood approach has clear advantages over the FIM approach in determining parameter confidence intervals, as it correctly predicts confidence intervals even for small amounts of data and non-linear models. Therefore, we suggest to use the profile-likelihood based confidence intervals as standard approach for mechanistic upstream process models. In addition to the exact determination of parameter confidence intervals, the profile-likelihood analysis is a guide to changes in the model structure that reduce non-identifiability and should lead to certain predictions that can be used for with high confidence in bioprocess development.

## Conclusion

We show in this study that profile-likelihood can be applied to determine parameter confidence intervals and non-identifiabilities of state-of-the-art macroscopic upstream process models. Currently parameter non-identifiabilities have been tackled by regularization (Anane et al., 2019; Ulonska et al., 2018) and Bayesian approaches (Hernández Rodríguez et al., 2019), here we instead adopt an approach where the model structure is adapted to alleviate non-identifiabilities guided by the profile likelihood. This approach has successfully been applied in systems biology (Maiwald et al., 2016) and has the advantage that no priors or regularization parameters need to be defined. However structural changes might decrease the model fit and then models need to be selected carefully.

The profile-likelihood approach gives true confidence intervals even with small amounts of data (i.e. a single run) and has therefore clear advantages over FIM-based approaches. Since parameter confidence intervals directly influence the confidence intervals of predictions, it is necessary to *exactly* determine those confidence intervals. When models are used for predictions that are relevant for e.g. the biologics license application (BLA) of regulatory authorities, parameter identifiability is a key step in the characterization of the risk associated with the model predictions. Hence, it is an essential prerequisite for the commercialization of modeling approaches.

## Acknowledgment

We would like to thank Alireza Ehsani, Sabine Arnold and Jens Smiatek for critical discussion of the results and Boehringer Ingelheim BPAD and IU for provision of the experimental data.

## Conflict of interest

The authors declare no conflict of interest.

## Funding

This research received no external funding.

## Nomenclature

c_glc_: glucose concentration [g/L]
c_gln_: glutamine concentration [g/L]
c_lac_: lactate concentration [g/L]
c_NH4_: ammonia concentration [g/L]
c_P_: product concentration [g/L]
c_target_: target glucose or glutamine concentration [g/L]
DCC: dead cell concentration [10^9^ cells/L]
F_feed_: feed rate [L/h]
k_d_: death rate [1/h]
k_d,max_: maximal death rate [1/h]
k_glc_: affinity constant to glucose concentration [g/L]
k_gln_: affinity constant to glutamine concentration [g/L]
k_lac_: affinity constant to lactate concentration [g/L]
k_lysis_: lysis rate of dead cells [1/h]
k_μ_: inhibitory effect of growth rate on death rate [1/h]
m_glc_: glucose maintenance [g/10^9^ cells*h]
q_glc_: specific glucose consumption rate [g/10^9^ cells*h]
q_glc,max_: maximal specific glucose uptake rate [g/10^9^ cells*h]
q_gln_: specific glutamine uptake rate [g/10^9^ cells*h]
q_gln,max_: maximal specific glutamine uptake rate [g/10^9^ cells*h]
q_lac_: net rate of specific lactate metabolism [g/10^9^ cells*h]
q_lac,con_: specific lactate consumption rate [g/10^9^ cells*h]
q_lac,max_: maximal specific lactate uptake rate [g/10^9^ cells*h]
q_lac,prod_: specific lactate production rate [g/10^9^ cells*h]
q_NH4_: specific ammonia production rate [g/10^9^ cells*h]
q_p_: specific production rate [g/10^9^ cells*h]
r_NH4,gln_: rate of glutamine decay [1/h]
VCC: viable cell concentration [10^9^ cells/L]
ʋ_lac,glc_: inhibitory effect of glucose consumption on lactate consumption [g/10^9^ cells*h]
ʋ_lac,gln_: inhibitory effect of glutamine consumption on lactate consumption [g/10^9^ cells*h]
ʋ_lac,prod_: inhibitory effect of lactate consumption on lactate production [g/10^9^ cells*h]
Y_BM/glc_: cells formed on glucose [10^9^ cells/g]
Y_BM/gln_: cells formed on glutamine [10^9^ cells/g]
Y_BM/lac_: cells formed on lactate [10^9^ cells/g]
Y_gln/dGG_: glutamine yield on media supplement [g/g]
Y_lac,glc_: lactate yield on glucose [g/g]
Y_NH4,gln_: ammonia yield on glutamine [g/g]
μ: growth rate [1/h]

## Notes

### Competing Interest Statement

The authors have declared no competing interest.

