## Supplemental Figures for "Reducing structural non-identifiabilities in upstream bioprocess models using profile-likelihood"

### Supplementary Nomenclature

$c_{bolus}$ – glucose or glutamine bolus concentration [g/L]

$c_{glc,bolus}$ – glucose concentration in bolus addition [g/L]

$c_{glc,feed}$ – glucose concentration in feed [g/L]

$c_{gln,bolus}$ – glutamine concentration in bolus addition [g/L]

D – dilution [1/h]

$F_{glc,bolus}$ – glucose bolus addition rate [L/h]

$F_{gln,bolus}$ – glutamine bolus addition rate [L/h]

LCC – lysed cell concentration [$10^9$ cells/L]

TCC – total cell concentration [$10^9$ cells/L]

$V_L$ – liquid volume [L]

### Adapted model from Ulonska et. al.

The model was adapted from Ulonska et. al.. The differential equation system was set as follows:

Viable cell concentration: $$\frac{dVCC}{dt} = \mu * VCC - k_d * VCC - VCC * D$$

Supplementary Equation 1

Dead cell concentration: $$\frac{dDCC}{dt} = k_d * VCC - k_{lysis} * DCC - DCC * D$$

Supplementary Equation 2

Lysed cell concentration: $$\frac{dLCC}{dt} = k_{lysis} * DCC - LCC * D$$

Supplementary Equation 3

Glucose concentration: $$\frac{dc_{glc}}{dt} = -q_{glc} * VCC + c_{glc,feed} * \frac{F_{feed}}{V} + c_{glc,bolus} * \frac{F_{glc,bolus}}{V} - c_{glc} * D$$

Supplementary Equation 4

Glutamine concentration: $$\frac{dc_{gln}}{dt} = -q_{gln} * VCC + Y_{\frac{gln}{dGG}} * r_{dGG} * c_{dGG} + c_{gln,bolus} * \frac{F_{gln,bolus}}{V} - r_{\frac{NH4}{gln}} * c_{gln} - c_{gln} * D$$

Supplementary Equation 5

Supplement concentration: $$\frac{dc_{dGG}}{dt} = -r_{dGG} * c_{dGG} - c_{dGG} * D$$

Supplementary Equation 6

Lactate concentration: $$\frac{dc_{lac}}{dt} = -q_{lac} * VCC - c_{lac} * D$$

Supplementary Equation 7

Ammonia concentration: $$\frac{dc_{NH4}}{dt} = -q_{NH4} * VCC + Y_{\frac{NH4}{gln}} * r_{\frac{NH4}{gln}} * c_{gln} - c_{NH4} * D$$

Supplementary Equation 8

Product concentration: $$\frac{dc_P}{dt} = q_P * VCC - c_P * D$$

*Supplementary Equation 9*

Volume: $$\frac{dV}{dt} = F_{feed} + F_{glcbolus} + F_{glnbolus}$$

*Supplementary Equation 10*

The kinetic expressions are given by:

Growth rate: $$\mu = Y_{\frac{BM}{glc}} * max\left(q_{glc} - \frac{q_{lac,prod}}{Y_{\frac{lac}{glc}}} - m_{glc}, 0\right) + Y_{\frac{BM}{gln}} * q_{gln}$$

*Supplementary Equation 11*

Death rate: $$k_d = k_{d,max} * \frac{k_\mu}{k_\mu + \mu}$$

*Supplementary Equation 12*

Lactate consumption and production:

$$q_{lac,con} = q_{lac,max} * \frac{c_{lac}}{k_{lac} + c_{lac}} * \frac{\nu_{lac,gln}}{\nu_{lac,gln} + q_{gln}}$$

$$q_{lac,prod} = Y_{\frac{lac}{glc}} * q_{glc} * \frac{\nu_{lac,prod}}{\nu_{lac,prod} + q_{lac,con}}$$

$$q_{lac} = -(q_{lac,prod} - q_{lac,con})$$

*Supplementary Equation 13*

Glucose consumption: $$q_{glc} = q_{glc,max} * \frac{c_{glc}}{k_{glc} + c_{glc}} * \frac{\nu_{\frac{lac}{glc}}}{\nu_{\frac{lac}{glc}} + q_{lac,con}}$$

*Supplementary Equation 14*

Glutamine consumption: $$q_{gln} = q_{gln,max} * \frac{c_{gln}}{k_{gln} + c_{gln}}$$

*Supplementary Equation 15*

Ammonia production: $$q_{NH4} = Y_{\frac{NH4}{gln}} * q_{gln}$$

*Supplementary Equation 16*

Dilution: $$D = \frac{F_{feed} + F_{glcbolus} + F_{glnbolus}}{V}$$

*Supplementary Equation 17*

### Supplementary Methods

Linear approximations of confidence intervals are derived from the hessian matrix H given by:

$$H = \nabla^T \nabla \chi^2 |_{\hat{\theta}_i}$$

*Supplementary Equation 18*

From which the covariance matrix C is calculated by

$$C = 2 * H^{-1}$$

*Supplementary Equation 19*

The parameter confidence intervals are then given by:

$$\sigma_i^{\pm} = \hat{\theta}_i \pm \sqrt{\chi^2(\alpha, df) * C_{ii}}$$

*Supplementary Equation 20*

### Supplementary Table 1

*Supplementary Table 1: Comparison of Profile-Likleihood and Approximate Confidence Intervals*

| | Optimal Parameter Value | Lower Bound PL | Upper Bound PL | Lower Bound Approximate | Upper Bound Approximate |
|---|---|---|---|---|---|
| Y_NH4gln | -1.34 | -1.40 | -1.27 | -1.40 | -1.27 |
| **Y_bmglc** | 3.22 | 1.93 | Inf | -22.13 | 28.57 |
| **Y_bmgln** | 1.96 | 1.70 | Inf | 1.57 | 2.34 |
| Y_glnGG | -0.94 | -1.47 | -0.69 | -1.29 | -0.60 |
| Y_lacglc | -0.18 | -0.24 | -0.11 | -0.34 | -0.02 |
| k_glc | -3.05 | -3.10 | -1.20 | -14.18 | 8.07 |
| k_gln | -0.48 | -0.88 | 0.10 | -0.87 | -0.09 |

| | | | | | |
|---|---|---|---|---|---|
| **k_lac** | 3.06 | -Inf | 3.06 | -66.43 | 72.55 |
| k_lys | -2.43 | -2.98 | -2.18 | -2.76 | -2.10 |
| k_mu | -3.05 | -3.18 | -2.94 | -3.18 | -2.93 |
| kdm | -1.93 | -2.00 | -1.84 | -2.02 | -1.83 |
| **m_glc** | -3.59 | -Inf | -3.58 | -32.99 | 25.80 |
| nu_lacglc | -2.58 | -2.79 | -2.36 | -3.24 | -1.91 |
| **nu_lacgln** | -3.54 | -Inf | -3.25 | -3.98 | -3.11 |
| **nu_lacprod** | -0.24 | -0.35 | Inf | -30.83 | 30.35 |
| q_glcm | -1.15 | -1.32 | -1.08 | -1.36 | -0.94 |
| **q_glnm** | -3.41 | -Inf | -3.06 | -3.84 | -2.98 |
| **q_lacm** | 0.68 | -Inf | Inf | -69.07 | 70.44 |
| q_p | -2.83 | -2.86 | -2.79 | -2.86 | -2.79 |
| r_NH4gln | -1.59 | -1.65 | -1.39 | -1.66 | -1.52 |
| r_dGG | -0.39 | -1.57 | 1.00 | -1.77 | 0.99 |
